# A community audit of error control in crosslinking mass spectrometry

**DOI:** 10.1101/2025.04.27.649519

**Authors:** Lutz Fischer, Bruno C. Amaral, Micha Johannes Birklbauer, James E. Bruce, Paulo C Carvalho, Robert J. Chalkley, Milan A Clasen, Viktoria Dorfer, Fabio C Gozzo, A.J.R Heck, Michael R. Hoopmann, Claudio Iacobucci, Andrew Keller, Lars Kolbowski, Swantje Lenz, Diogo Borges Lima, Fan Liu, Karl Mechtler, Robert L. Moritz, Fränze Müller, Julia Ruta, David C. Schriemer, David D. Shteynberg, Michael J. Trnka, Rosa Viner, Juri Rappsilber

## Abstract

In a community effort, we audited error control in twelve crosslinking mass spectrometry pipelines using two ground-truth designs that expose a lower bound on false identifications. Some pipelines passed, whereas for others these verifiably false identifications alone exceeded the requested false discovery rate up to 29-fold at the protein–protein interaction level. The audit also tracked improvement between software versions, supporting its use for regression testing.

---

Crosslinking mass spectrometry (MS) converts spectra of covalently linked peptide pairs into residue distance restraints and protein–protein interactions (PPIs), making error control consequential at several reporting levels ^1^. Most pipelines estimate false discovery rates (FDRs) from matches to decoy sequences. This estimate is informative only if decoys remain representative of false target matches after candidate selection, scoring, rescoring, post-processing and aggregation. Crosslinking MS challenges this requirement because paired-peptide search spaces are large, pipeline-specific prefilters are common, and errors accumulate when crosslinked peptide-spectrum matches (CSMs) are collapsed into residue pairs or PPIs ^1–3^. A multi-lab comparison of complete workflows on a single protein ^4^ and a community call for agreed error assessment and benchmarking ^5^ preceded this effort. Here, twelve widely used pipelines, mostly run by their developers, analysed identical crosslinking data from lysates and whole cells.

Entrapment databases, synthetic standards, and mixed-species experiments have already shown that nominal crosslinking MS FDRs can underestimate empirical error ^6–11^. However, synthetic standards do not reflect the complexity, noise or dynamic range of current crosslinking experiments ^6,8,10^. In this community effort, the same crosslinking data from lysates and whole cells were analysed with twelve widely used pipelines, mostly by their developers. Two entrapment designs we developed previously ^7,9^ provide partial ground truth. Because they reveal only a subset of false identifications, termed observable false targets (OFTs), the OFT rate (OFTR) is a lower bound on the true error and should not exceed a correctly estimated FDR (Fig. 1a). In addition, comparing the score distributions and numbers of decoys and OFTs tests whether decoys model the observable error (Fig. 1b).

**Fig. 1.**
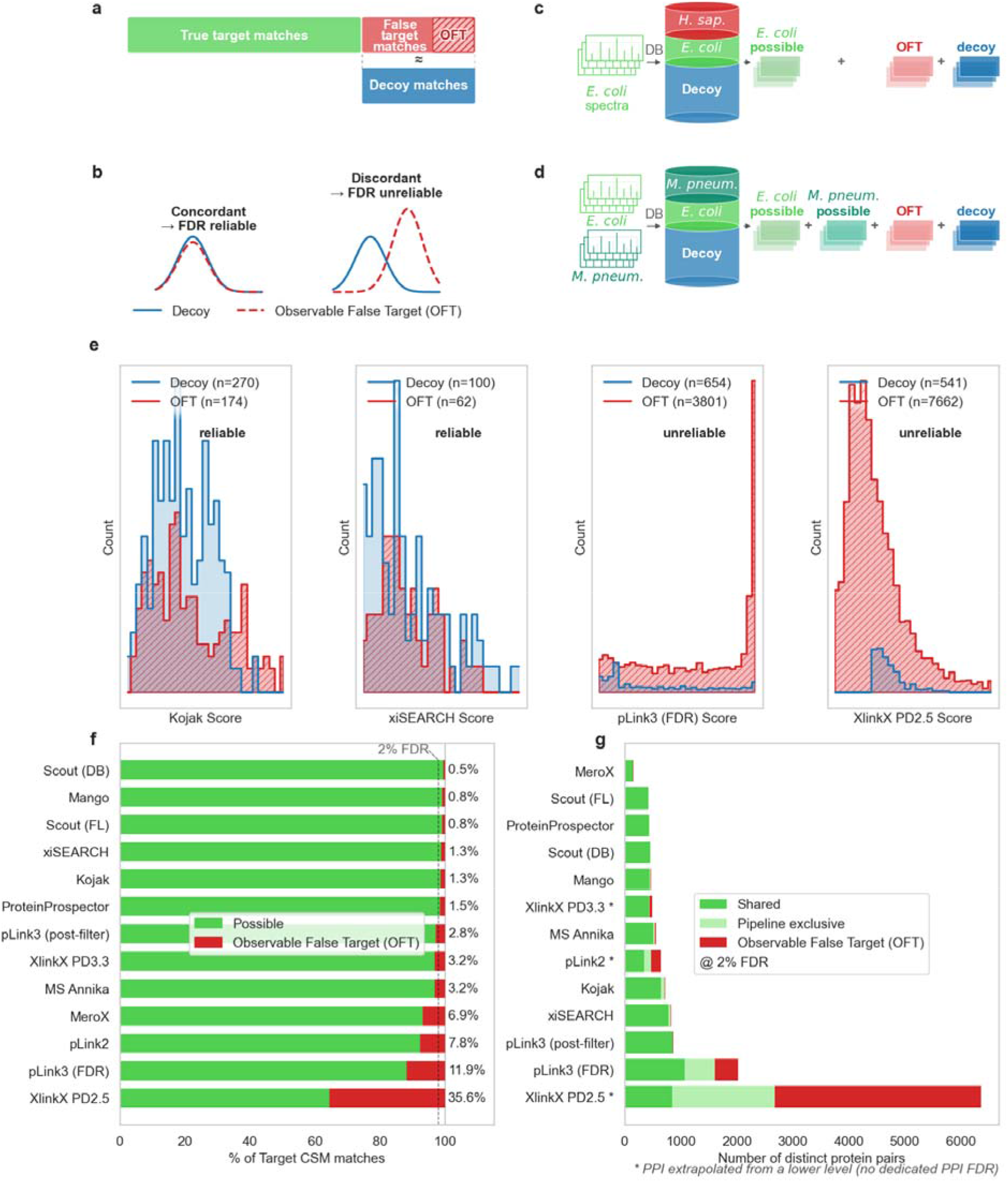
Decoys and observable false targets across crosslinking MS pipelines. **(a)** Target–target CSMs are either true (green) or false (red); a subset of false positives is observable false targets (OFT, hatched). Decoys (blue) should model the false positives. **(b)** Concordant engines (left) show aligned decoy and observable FP score distributions with expected ratios; discordant engines (right) do not. **(c)** *E. coli*–*H. sapiens* design: the search database adds human proteins as entrapment, plus a decoy database of equal size, to the *E. coli* target; outcomes split into possible (could be correct, could be wrong), OFT (a human match, definitively false), and decoy (with the exception of Mango, which does not use decoys). **(d)** *E. coli*–*M. pneumoniae* design: spectra from each organism are searched against the combined target database of both species plus an equal-sized decoy database; outcomes split into possible (own-species match, could be correct, could be wrong), OFT (cross-organism match, definitively false), and decoy (except Mango). **(e)** Decoy (blue) and OFT (red, hatched) score distributions for four engines (*E. coli*–*H. sapiens*, CSM level) **(f)** Fraction of target CSMs classified as possible (green) or OFTs (lilac/red) per engine; ranked by increasing impossible fraction. **(g)** Distinct protein pairs per engine, split by whether they are shared across engines (dark green), exclusive to one engine (light green), or OFT due to one protein being exclusively *H. sapiens*.

We based our investigation on two complementary DSSO datasets covering *Escherichia coli* and *Mycoplasma pneumoniae*. As a single entrapment control, the *E. coli* data were searched against a combined *E. coli*–*H. sapiens* database. Any identification involving a human (entrapment) protein was an OFT (Fig. 1c). The number of entrapment matches should be at most half the number of decoy matches (Online Methods). As a dual entrapment control, data from separate *E. coli* and *M. pneumoniae* acquisitions were pooled and searched against an *E. coli*–*M. pneumoniae* database (Fig. 1d). Matches of a spectrum to the other species, and any cross-species protein pair, were therefore OFTs ^7,9^. The decoy-free Mango workflow contributed only to the lower-bound audit. Analyses were restricted to heteromeric crosslinks, which report on PPIs.

Several pipelines, for example Kojak and xiSEARCH, passed both tests: decoys and OFTs had similarly shaped score distributions at CSM level, and the OFTR stayed within the requested FDR (Fig. 1e; all pipelines in Supplementary Figs. 1 and 2). By contrast, for MeroX, both pLink versions and XlinkX PD2.5, OFTs outnumbered decoys 2- to 14-fold, although they should number at most half as many, and for pLink and XlinkX PD2.5 they also occupied score ranges depleted of decoys (Fig. 1e; Supplementary Fig. 1). At a requested FDR of 2%, OFTRs reached 6.9% for MeroX, 7.8% for pLink2, 11.9% for pLink3 after FDR and 35.6% for XlinkX PD2.5 in the *E. coli*–*H. sapiens* dataset (Figs. 1f and 2a,b; dual entrapment in Supplementary Figs. 3 and 4). pLink3 employs an additional post-FDR filtering step that reduces the OFTR for CSM to 2.8%, but the filter lies outside the error model, so its effect on other datasets or requested FDRs is unknown.

**Fig. 2.**
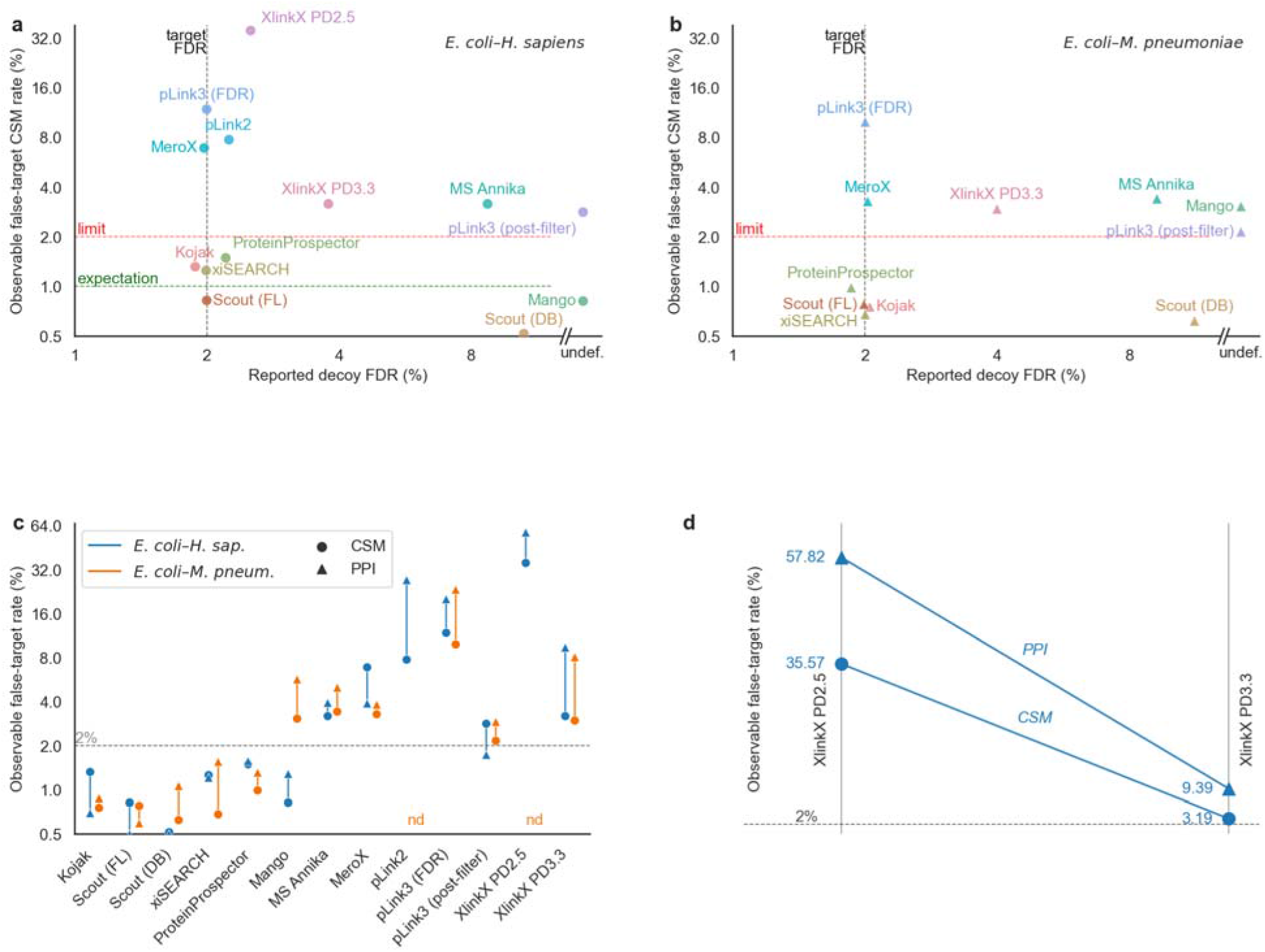
Consequences of decoy–false-target discordance. **a, b,** Declared decoy-based FDR (x-axis) vs. directly observable false-target CSM rate (y-axis) for each engine, on the *E. coli*–*H. sapiens* (**a**) and *E. coli*–*M. pneumoniae* (**b**) datasets. Dashed red and black lines mark 2% on both axes as target for the decoy FDR (black) and limit what the number of observable false positives should not exceed (red), green dashed line is the expectation for observable FP, if no match external information would have been used ; “undef.” indicates engines without a computable decoy FDR at CSM level. **c,** Observable false-target rate at CSM (circle) and PPI (triangle) level for each engine. Blue: *E. coli*–*H. sapiens*; orange: *E. coli*–*M. pneumoniae*. Engines with a large CSM-to-PPI increase export disproportionate numbers of false PPIs. Dashed line marks 2%. **d,** CSM- and PPI-level observable false-target rates for XlinkX PD2.5 and XlinkX PD3.3 (*E. coli*–*H. sapiens*). The substantial reduction between versions demonstrates the impact of community driven benchmarks. Dashed line marks 2%

These data distinguish low observable error in line with the requested FDR, elevated error signalled by decoys, and elevated error concealed by decoy–OFT discordance. The audit cannot reveal algorithmic causes, but the patterns point to possible mechanisms. For MS Annika, decoys modelled the error, but the FDR was not controlled separately for self-links and heteromeric links, leaving heteromeric links with higher error than requested ^7^. Mango and Scout (DB) showed markedly higher OFTRs with dual than with single entrapment, most clearly at the PPI level (Supplementary Fig. 3b,d). This suggests that these pipelines use information beyond the individual CSM in ways their error models do not capture ^9,12^.

The consequences were largest at the PPI level, where false matches accumulated as pipeline-specific protein pairs (Figs. 1g and 2c, Supplementary Fig. 5) ^3,7^. PPI-OFTRs were low for pipelines with dedicated interaction-level error control but rose sharply for several pipelines lacking it, reaching 57.8% for XlinkX PD2.5. These results guided a revision of XlinkX: XlinkX PD3.3 lowered the OFTR from 35.6% to 3.2% for CSMs and from 57.8% to 9.4% for PPIs (Fig. 2d). Much of the remaining error likely reflects the PPI-level FDR that XlinkX has yet to implement.

Our audit shows that FDRs reported by crosslinking MS pipelines cannot be taken at face value. Observable false targets bound the error only from below, so they cannot certify a pipeline, but they can expose one whose decoys no longer represent its false matches. The true error can be extrapolated from the OFTR in some, but not all, cases (Online Methods). Encouragingly, several pipelines combined low observable error with dedicated PPI-level error control, indicating that reliable interaction maps are within reach when error is controlled at the level used for biological interpretation. As the revision of XlinkX shows, the audit doubles as a regression test during software development. We provide the data and code so that such audits can become routine for developers and users alike. Extending such audits to other cleavable and non-cleavable crosslinkers, photo-crosslinkers and enrichable crosslinkers, and to further datasets, is a natural next step.

## Methods

### Datasets

#### E. coli–H. sapiens (ecoli_human)

Prior to crosslinking, the *E. coli* K-12 lysate was fractionated by size-exclusion chromatography (SEC) into 44 fractions (ECLP_ID_01 - ECLP_ID_44). Each fraction was crosslinked individually with DSSO (Disuccinimidylsulfoxide) before fractions were pooled, trypsin-digested, and re-fractionated for MS acquisition. A protein was considered detectable as part of a crosslink in a fraction when its iBAQ value exceeded 1 × 10□. Protein intensities per fraction were quantified by label-free proteomics and are publicly available at JPST000843 / PXD019004. Two *E. coli* proteins that were never co-detected in any fraction above this threshold are assumed to not be observable as part of a crosslink. The combined database included *E. coli* proteins and human proteins added as entrapment. Raw mass spectrometry data: jPOST JPST000845 / PXD019120 ^7^.

#### E. coli–M. pneumoniae (ecoli_mycpn)

*E. coli* K-12 and Mycoplasma pneumoniae lysates were prepared, DSSO-crosslinked, and trypsin-digested, and the spectra independently acquired. Their runs were concatenated into a single combined search against a combined E. coli + Mycoplasma pneumoniae database. The originating organism of each scan was recorded from scan-number split points stored per engine in input/merged_files.json, constructed at analysis time from the constituent single-species run assignments. Raw data: jPOST JPST000845 / PXD019120, PRIDE PXD017711 ^13^.

### Observable False Target classification

An identification is classified as observable false target (OFT) when it violates a constraint fixed by the experiment, evaluated only among target–target (TT) matches.

#### Single entrapment - absent protein and non-co-present pairs (*E. coli H. sapiens*)

A crosslink is an OFT when at least one of the peptides is unambiguously from a human or decoy protein. Incorrect matches can come from both *E. coli* and *H. sapiens* databases but as known false positives (FP) we can only count target *H. sapiens* matches. Among false positives we would predominantly expect false matches where only one peptide is incorrectly identified and we would expect this incorrect peptide to be from a decoy sequence twice as often as from a sequence from the target database. If the Observed FP were noticeably lower than the expected ratio this would be an indication that match-external information is used somewhere in the pipeline (e.g. scoring or filtering). Match-external here means additional information independent of the spectrum and the matched peptide, for example the information that a protein was observed with self links or hydrolysed crosslinker modification. As the data were acquired with protein level fractionation before crosslinking ^7^, a pair of *E. coli* proteins can also be considered an OFT crosslink if there is no SEC fraction in which both proteins have an iBAQ value ≥ 1 × 10⁶. (Sup. Fig S3)

#### Dual entrapment - per-spectrum species origin (ecoli_mycpn)

Both proteomes are genuinely present in the combined sample, so an OFT is not defined by absence but by the spectrum-conditioned observability of each protein. Each spectrum’s true organism is known from its run of origin; for that spectrum, only proteins of its origin species are observable. A crosslink is therefore an OFT if the species assignment of at least one of its peptides conflicts with the spectrum’s recorded origin, which also makes every cross-proteome (*E. coli*–*M. pneumoniae*) link an OFT. Because the impossible partners are present, plausible proteins rather than absent ones, this classification additionally detects matches that a pipeline has up-weighted using protein or peptide presence or crosslinkability information not represented in its error model. If a search pipeline performs noticeably better in the single entrapment case than in the dual entrapment case then this is a good indication that match-external information is used and possibly, if the detectable error exceeds the stated error, not correctly considered during error modeling (Sup. Fig S4). The OFTR is the fraction of TT crosslinks that are impossible.

At the PPI level, the same definitions are applied to reported protein pairs rather than spectral matches. However, for most search engines, the individual PPIs cannot be traced back to their supporting spectrum matches. Even if this information were available, determining whether a within-species PPI should be considered an OFT would not be straightforward if it were supported by spectra originating from both species. We therefore restrict the OFTs at the PPI level to cross-proteome (E. coli–M. pneumoniae) pairs, which are impossible under the per-spectrum species-origin constraint.

### The observable false target rate as a lower bound

OFT matches are a strict subset of all false positives: a false crosslink between two genuinely present proteins of the correct species is not detectable as impossible. Consequently the OFTR never exceeds the true FDR (OFTR ≤ FDR), and an OFTR above a pipeline’s declared FDR proves miscalibration with no further assumption.

The bound can optionally be scaled to an approximate point estimate. If f denotes the fraction of false matches that are detectable as impossible, set by database composition (e.g. the relative sizes of the present and absent, or the two species’, databases), then true FDR ≈ OFTR / f. For the single entrapment dataset f would be ∼0.5 as the entrapment proteins were selected to resemble target proteins. However, for the dual entrapment dataset combined whole proteome FASTAs were used and the effective value of f is then different for CSM and PPI. While this is still possible to consider, post search processing, e.g. by considering match external information, changes the effective database size. This change is not known to us and thus impossible to consider, making correct scaling impossible. The second dataset is meant to test for the effect on FDR estimations of also such database changes during post-processing.

### Normalisation pipeline

Per-pipeline loader functions map raw search output to a canonical schema, normalising decoy and entrapment Accession number. Protein pairs are classified as protein heteromeric or self link by stripping decoy prefix from accession numbers and checking the overlap of the possible accessions for peptide one and two. Loader output is concatenated with a search_engine column for each dataset and level. For ecoli_mycpn the spectrum origin is added and a flag is set if at least one possible protein source for each peptide matches the spectrum origin species.

### Cross-pipeline pair overlap

Peptide pairs (CSM level) and protein pairs (PPI level) are compared across pipelines using a canonical, direction-invariant key, so that a crosslink reported with its two peptides or proteins in either order counts as the same pair. Within one engine, repeated identifications of the same pair - across different spectra, charge states, or modification states - collapse to a single distinct pair rather than being counted once per spectrum match. Peptide sequences are canonicalised before keying to remove notation that differs by engine but does not change which peptide was identified: bracketed or engine-specific modification tags are stripped, isoleucine and leucine are treated as equivalent (as pLink does so), and a leading methionine on n-terminal peptides is removed. This protein’s initiator methionine is removed, as some pipelines include it in the reported sequence and others omit. Protein accessions are canonicalised by sorting ambiguous accession groups and stripping the entrapment tag, so the same protein is recognised whether or not a pipeline flags it as entrapment. Overlap is only counted on target-target matches. A pair is counted as shared when reported by more than one pipeline and unique when reported by only one; the possible/OFT split for each pair follows the impossible-match classification above (Fig. 1g, Fig. S5, Fig S6).

### Declared (decoy) FDR

The decoy-based FDR is recalculated from the same normalised output and treated as the pipeline’s reported error estimate - the quantity being audited - rather than as an independent estimator. Crosslinks are classified into TT, TD (one decoy), and DD (both decoy) using Decoy1/Decoy2, and decoy-based FDR is calculated as ^2,3,14^:

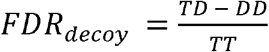

For XlinkX PD2.5, whose DD count >> TD count This would lead to negative FDRs, hence we changed it to a non-standard decoy handling, (TD + DD) / TT; this decoy estimate is reported for completeness but this is probably more of an expression of underlying problems than an actual meaningful FDR calculation. For any given FDR TD should be larger than DD as even for completely random data the TD space is twice as large as the DD space and TD not only contains matches with both peptides wrong but also cases with only one peptide wrong and one correctly identified. Hence TD should at least be twice DD.

### Search engine configurations

Twelve crosslink search pipelines were benchmarked: Kojak ^15,16^ (v2.0.2; PPI-level scoring via iProphet ^17^ (using Trans-Proteomic Pipeline v7.1.0 Albedo)), Scout ^10^ (Version 2.0.0, and Version 2.1.0), xiSEARCH ^18^ (v1.8.13), ProteinProspector ^19^ (v6.5.2) with subsequent rescoring, summarisation and FDR control performed using a pre-release version of Touchstone ^20^, MS Annika ^21^ (v3.0.7), Proteome Discoverer (v3.1.0.638), MeroX ^22^ (v2.0.2.5), Mango ^23^ (decoy-free, v20260105), pLink2 ^24^ (v2.3.10, run in-house), pLink3 ^24^ (v3.0.17, third-party submitted results), XlinkX PD2.5 ^25^ (Proteome Discoverer 2.5), and XlinkX PD3.3 ^26^ (Proteome Discoverer 3.3.0.386). Scout is represented by two independently developed versions, Scout (FL) and Scout (DB), which have diverged from a common ancestor and differ in their search settings (below). They are counted and reported as separate pipelines, as are the two pLink versions and the two XlinkX versions.

All searches used DSSO as the crosslinker and trypsin as the protease, with carbamidomethylation of cysteine as a fixed and oxidation of methionine as a variable modification. Precursor and fragment mass tolerances were harmonised across pipelines at 4 ppm and 8 ppm respectively; this is confirmed in every parameter file we received. A 2% FDR was requested in every submission. Remaining settings followed each pipeline’s defaults for DSSO searches, with the following deviations:

- Missed cleavages were limited to two in all submissions except Scout (DB), which allowed three.
- The residues carrying the crosslinker differed. MS Annika, XlinkX, ProteinProspector and Scout (FL) searched DSSO on lysine and the protein N-terminus; xiSEARCH additionally allowed serine, threonine and tyrosine at reduced weight; Scout (DB) searched lysine, serine, tyrosine and threonine with equal weight.
- Peptide length limits differed, most markedly for MeroX, which considered peptides of 7–23 residues against 5–60 or wider for the other pipelines. Scout (FL) used a minimum peptide length of 7.
- In the XlinkX workflow the harmonised tolerances apply to the crosslink search node; an upstream Sequest HT search (20 ppm precursor and 0.5 Da fragment for PD2.5 and 10ppm and 0.02 Da for PD3.3) and a spectrum confidence filter precede crosslink detection.
- ProteinProspector generated its decoy database by randomisation rather than reversal. Touchstone parameters were minPepLen = 5, minProdIon.1 = minProdIon.2 = 4, and decoyScalingFactor = 1. For a short overview of used parameters see supplemental table S5.

Where a pipeline did not report PPI-level error control natively, PPI-level information was generated by extracting unique protein pairs from the lower level without making any additional cut-offs.

### Code and data availability

Normalisation pipeline, loaders, analysis scripts, and figure-generation code: https://github.com/Rappsilber-Laboratory/SingleDualEntrapment. Python 3.10+; dependencies pinned in Pipfile.lock (pandas, pyarrow, matplotlib, seaborn, pytest). Raw crosslinking MS data: jPOST JPST000845 / PXD019120. Normalised combined.parquet files, the per-dataset and detectable fractions: PRIDE [upload in progress].

Touchstone source code and documentation are available at https://github.com/mtrnka/Touchstone. The analysis reported here used a pre-release development version.

## Acknowledgements

We thank Timo Sachsenberg and Oliver Kohlbacher for running OpenPepXL in an earlier version of this study. This work was funded by the Deutsche Forschungsgemeinschaft (DFG, German Research Foundation) under Germany’s Excellence Strategy – EXC 2008 – 390540038 – UniSysCat, by the European Research Council (ERC) under the European Union’s Horizon Europe research and innovation programme (grant agreement No. 101119142 – TransFORM), in part by the National Institutes of Health grants R01 GM087221 (R.L.M.) and S10 OD026936 (R.L.M.), and by the National Science Foundation grant MRI-1920268 (R.L.M.).

## Author contributions

L.F. and J.R. conceived and designed the study. L.F. developed the analysis pipeline and performed the comparative analyses. S.L. and L.K. contributed to the initial study and to the development of xiSEARCH and xiFDR. Search results were provided by: L.F. (xiSEARCH); L.K. (pLink2, XlinkX PD2.5); M.R.H., D.D.S. and R.L.M. (Kojak, Trans-Proteomic Pipeline); J.E.B. and A.K. (Mango); R.J.C. and M.J.T. (ProteinProspector, Touchstone); M.J.B., V.D., F.M. and K.M. (MS Annika); F.L. and J.Ru. (Scout (FL)); M.A.C., D.B.L., P.C.C. and F.C.G. (Scout (DB)); C.I. (MeroX); B.C.A. and D.C.S. (pLink3); R.V. (XlinkX PD3.3). J.R. coordinated the community effort. L.F. and J.R. wrote the manuscript with input from all authors.

---

**Figure S1.**
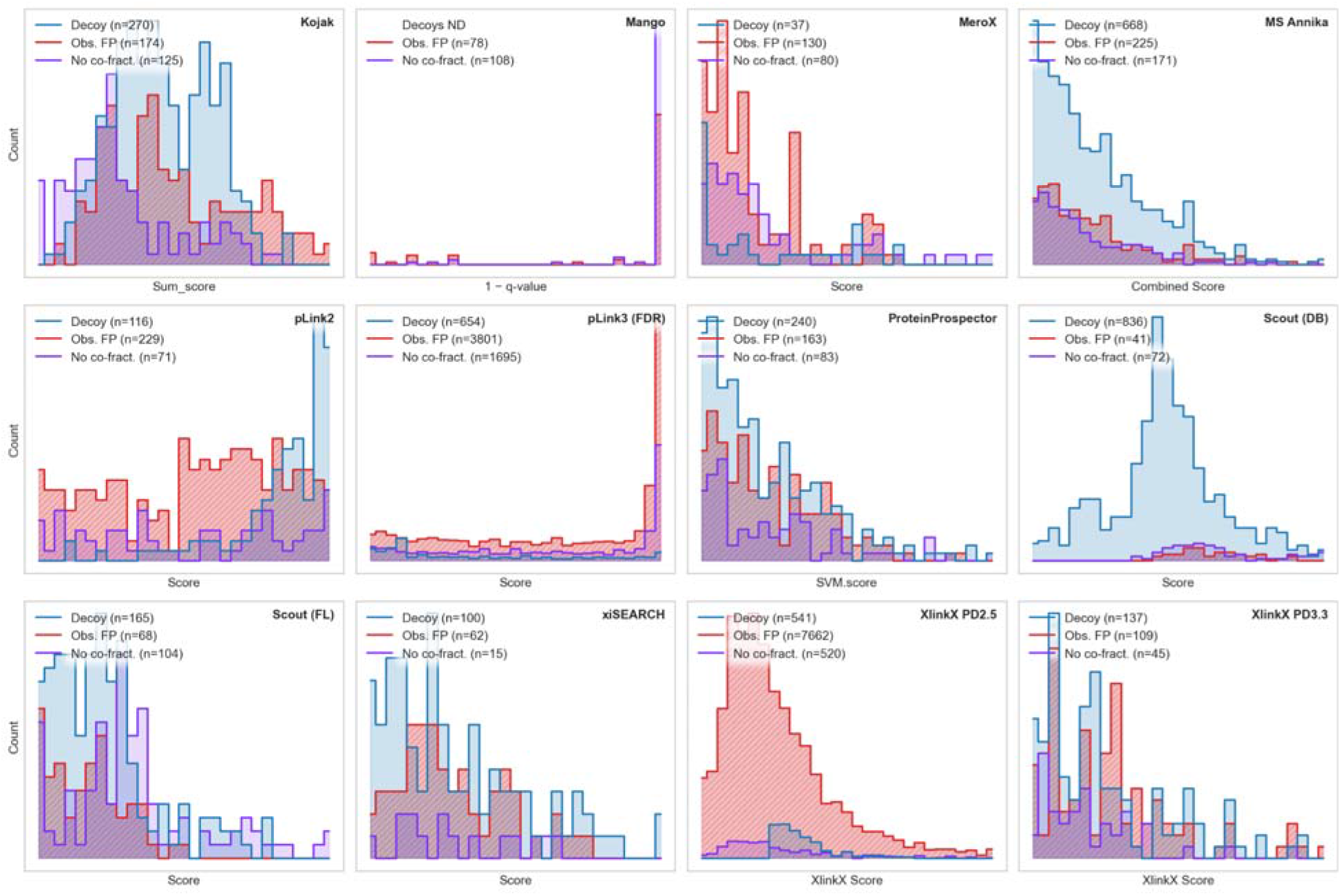
Score Distributions of Observable False Positives and Decoys in *E. coli*–*H. sapiens* dataset. One panel per pipeline, showing the score distribution of decoy matches (blue), of TT matches to a human entrapment protein (red) and of TT *E. coli*–*E. coli* pairs whose proteins never co-fractionate (purple). The three classes are disjoint. If decoys model false target matches correctly, the blue and red distributions should have the same shape and the blue count should be about twice the red. Pipelines where the red distribution instead extends to high scores, or outnumbers the decoys, have a decoy population that no longer represents their error.

**Figure S2.**
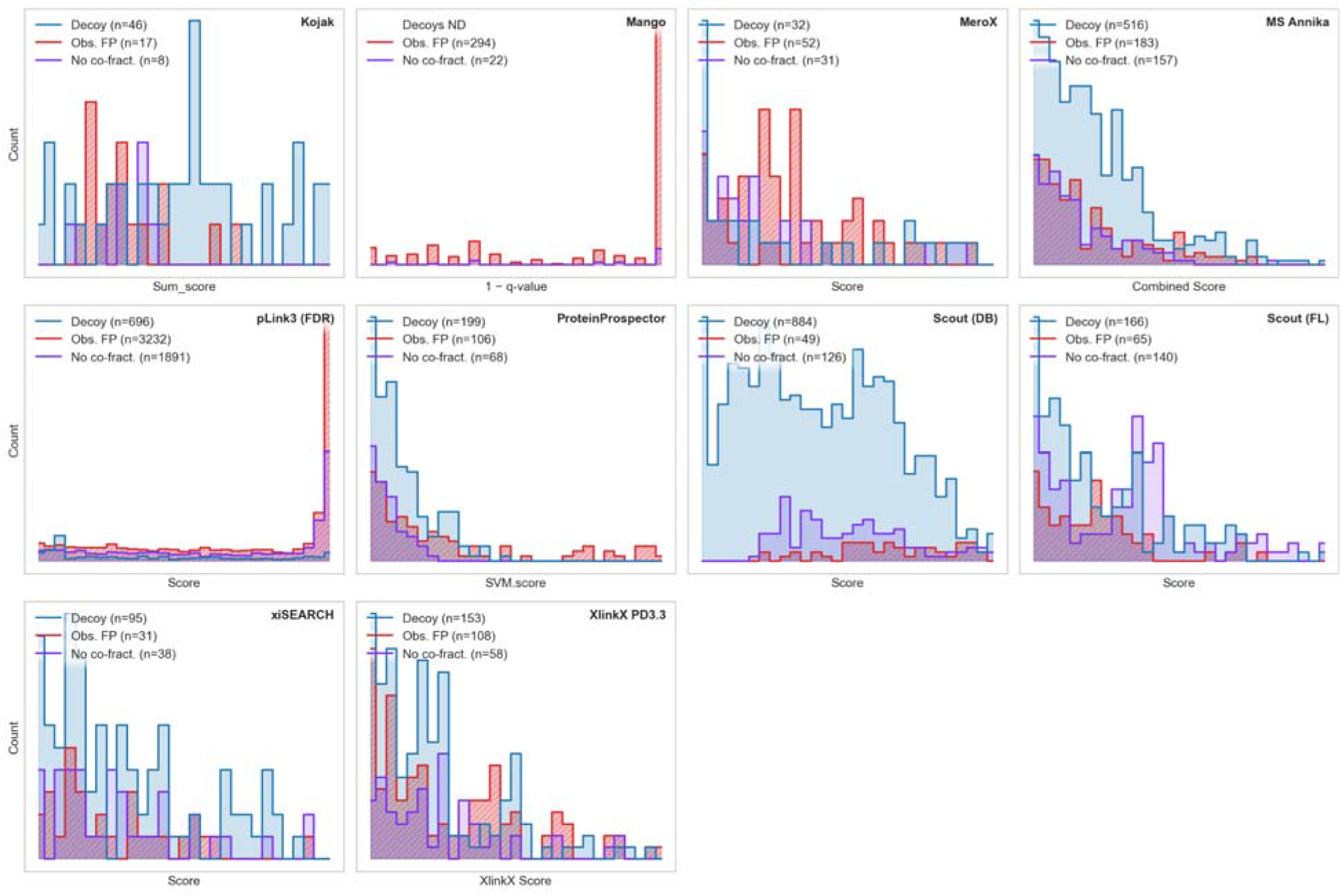
Decoy vs known false-positive score distributions (*E. coli*–*M. pneumoniae*, CSM) The *E. coli*–*M. pneumoniae* counterpart of Figure S1, with the ground truth supplied by the other reference dataset: a CSM whose spectrum came from one organism but was matched to the other (red). Decoy matches are blue; *E. coli*–*E. coli* pairs that never co-fractionate are purple. Reading the two figures side by side separates pipeline behaviour from any quirk of a single ground-truth construction. Note that the expected decoy-to-false-positive ratio is not 2 here: only the cross-species subset of false positives is observable, and the two organisms contribute very unequal shares of the database.

**Figure S3.**
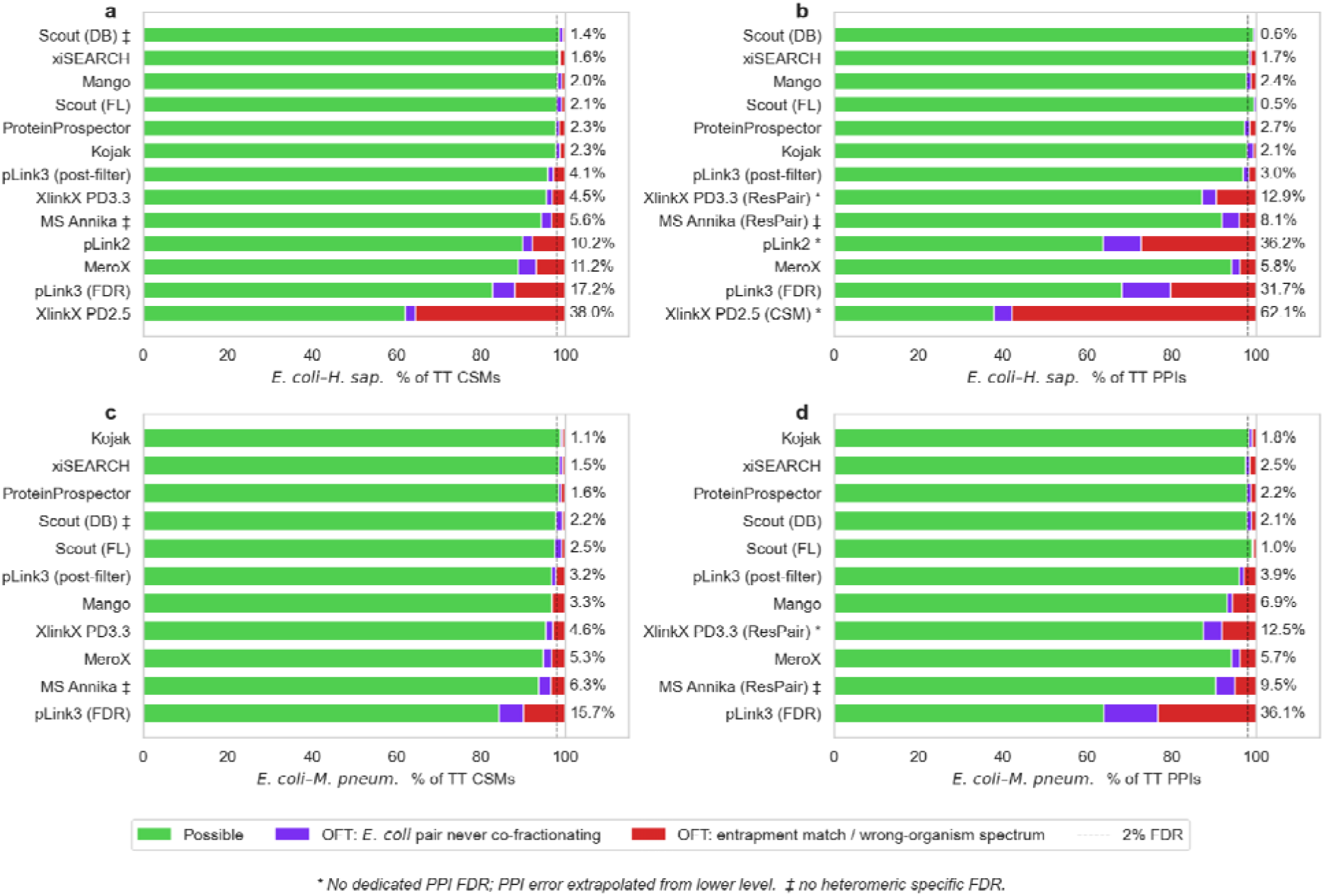
Observable false target (OFT) breakdown. Composition of the accepted target–target matches for each pipeline, as a percentage of TT. **(a, b)** *E. coli*–*H. sapiens* at CSM and PPI level. **(c, d)** *E. coli*–*M. pneumoniae* at CSM and PPI level. Bars run possible (green), then *E. coli* pairs that never co-fractionate (purple), then entrapment matches or wrong-organism spectra (red). The number at the right of each bar is the total OFTR. Each dataset is ranked on its own CSM total, and its PPI panel repeats that order so a pipeline sits at the same height in both. Pipelines marked * have no dedicated PPI FDR, with PPI error extrapolated from a lower level; ‡ marks pipelines that do not split FDR by link type.

**Figure S4.**
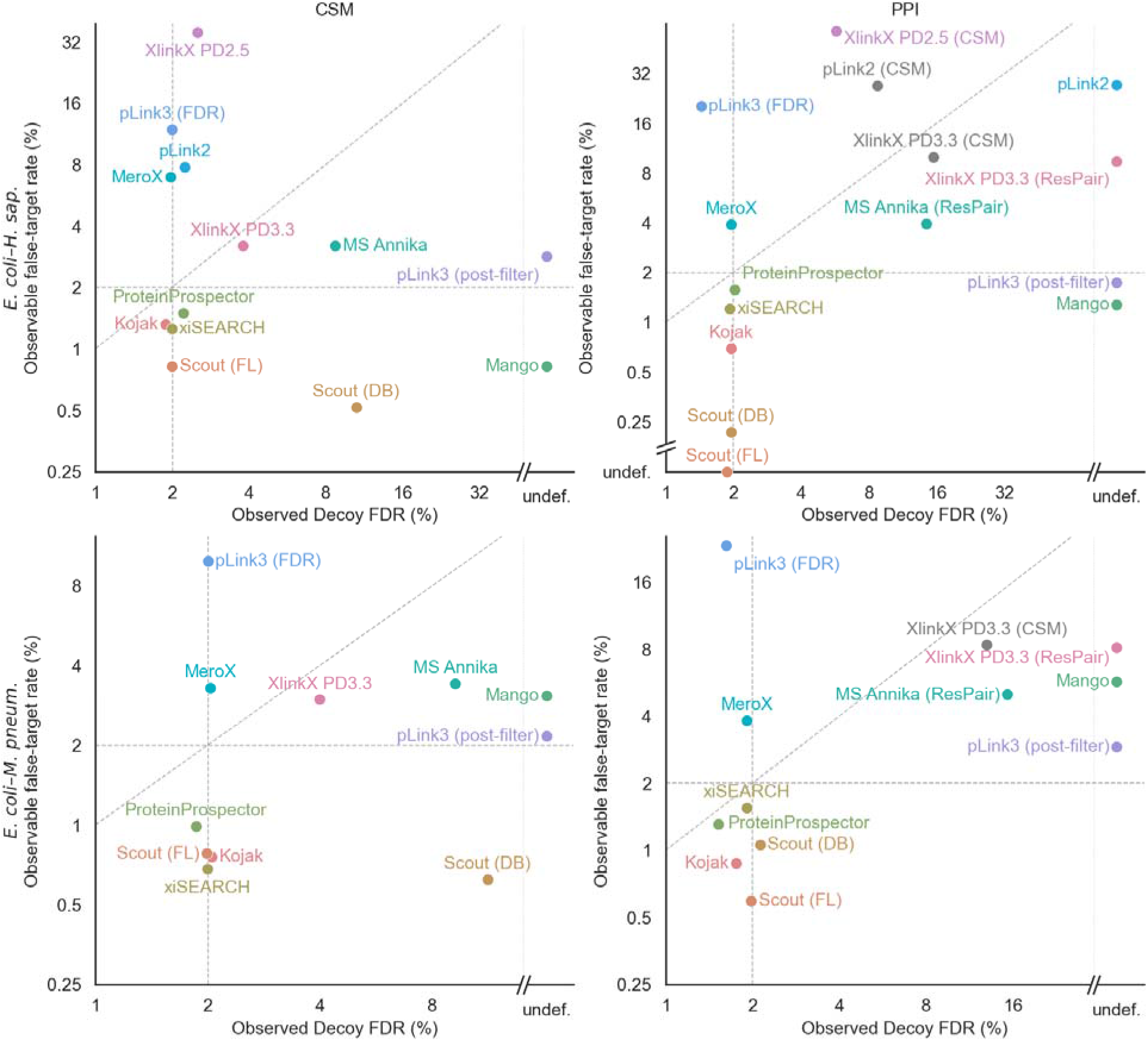
Observable error vs observed decoy FDR. Each point is one search pipeline at its declared 2 % FDR; x is the observed decoy FDR, y the OFTR. Rows are the two datasets (*E. coli*–*H. sapiens* top, *E. coli*–*M. pneumoniae* bottom), columns the level (CSM left, PPI right). Points above the diagonal under-report their error. Mango, which does not use decoys for error estimate, and PPI level information without supplied decoys are placed in a separate column at the right. Scout (FL) *E. coli*–*H. sapiens* did not produce any detectable false positives and was put on the x-axis. These are marked as undefined.

**Figure S5.**
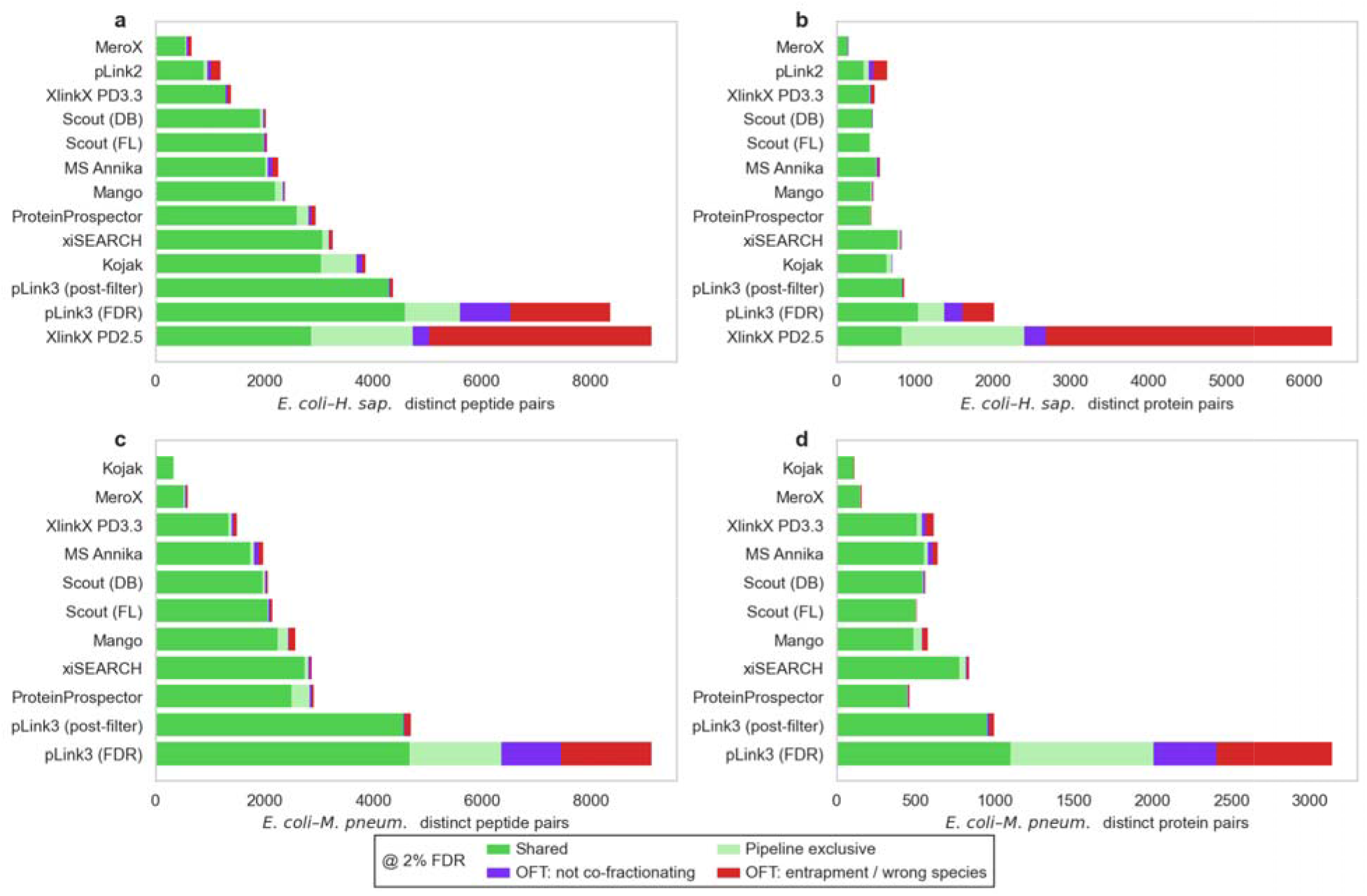
Cross-pipeline overlap. Overlap of reported pairs across pipelines, for both datasets and both levels. Rows are the datasets (*E. coli*–*H. sapiens* top, *E. coli*–*M. pneumoniae* bottom); columns are the level (unique peptide pairs left, unique protein pairs right). Bars split into shared vs unique and possible vs observable false positives (FP). Observable FP are again split into either an entrapment or cross-species match, or an *E. coli* pair whose two proteins never co-fractionated. Each row shares one engine order, taken from its peptide-pair panel, so a pipeline sits at the same height in both of its panels.

**Figure S6.**
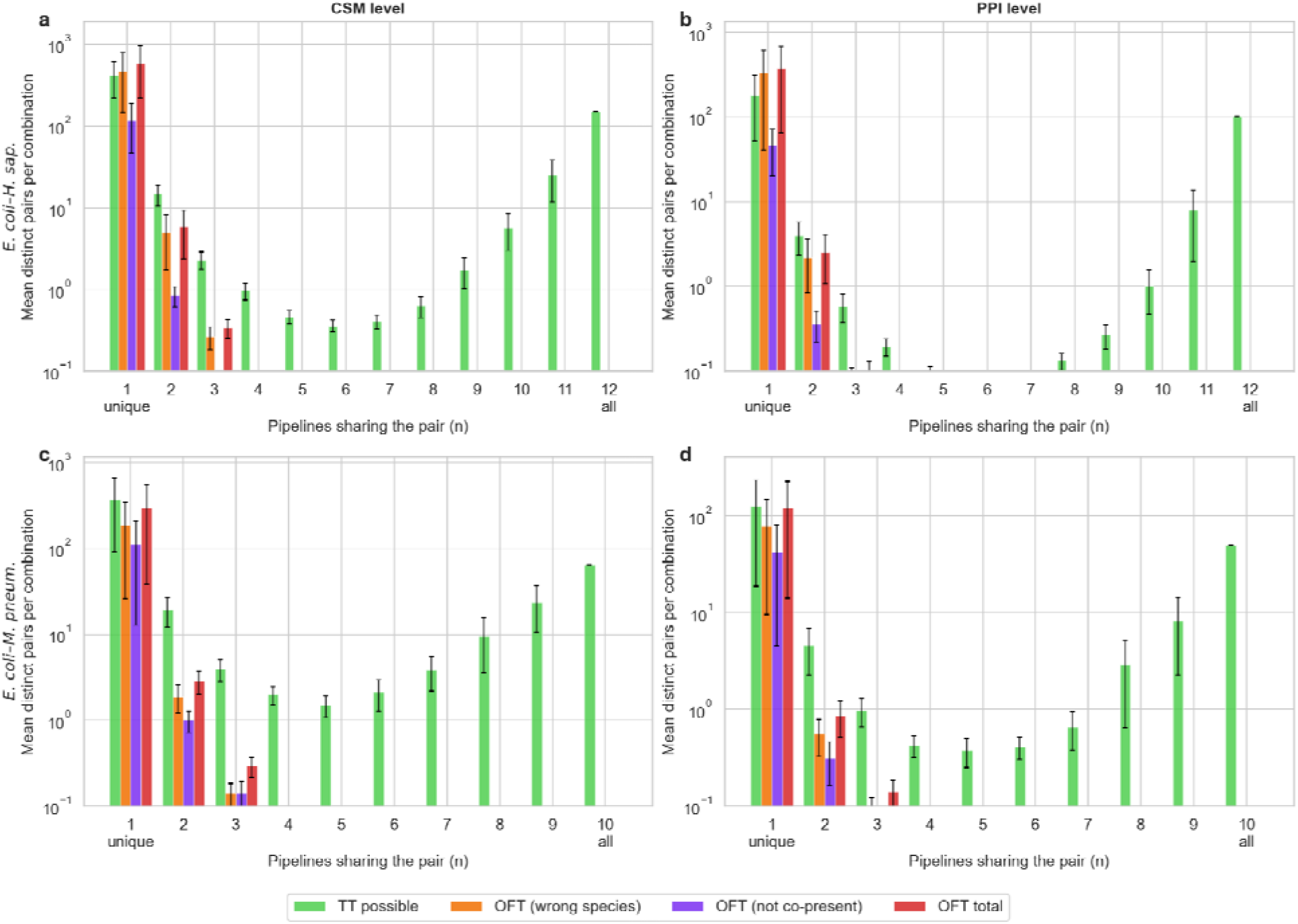
Cross pipeline overlap by pipeline count. Extension of Figure S5 (cross-pipeline overlap): instead of one shared/unique split per pipeline, pairs are grouped by how many of the N pipelines report them (pLink3 post-filter excluded; N = 12 for E. coli–H. sapiens, N = 10 for E. coli–M. pneumoniae). Rows are the datasets (E. coli–H. sapiens top, E. coli–M. pneumoniae bottom); columns are the level (CSM left, PPI right) — the same layout as Figure S5. (a) E. coli–H. sapiens, CSM level. (b) E. coli–H. sapiens, PPI level. © E. coli–M. pneumoniae, CSM level. (d) E. coli–M. pneumoniae, PPI level. For each sharing level n = 1 (unique to one pipeline) through n = N (reported by every pipeline), bars give the mean distinct-pair count across all C(N, n) specific n-pipeline combinations — not just the combinations that happen to occur — split into TT possible (green), observable false target (OFT) from a wrong-species/entrapment match (orange), OFT from an E. coli pair that never co-fractionates (purple), and the two OFT reasons pooled (OFT total, red). Error bars are the standard error across combinations; they are absent at n = N, where only one such combination exists. The y-axis is log-scaled, so a category with zero pairs at a given n is simply not drawn.

**Supplemental Table S1.** Single entrapment CSM level data per search engine.

| Search Pipeline | TT | TD | DD | Decoy FDR | OFT Species | OFTR Species | OFT not copresent | OFTR combined |
| --- | --- | --- | --- | --- | --- | --- | --- | --- |
| Kojak | 13204 | 259 | 11 | 1.88% | 174 | 1.32% | 125 | 2.26% |
| Scout (FL) | 8268 | 165 | 0 | 2.00% | 68 | 0.82% | 104 | 2.08% |
| Scout (DB) | 7919 | 834 | 2 | 10.51% | 41 | 0.52% | 72 | 1.43% |
| xiSEARCH | 4931 | 99 | 1 | 1.99% | 62 | 1.26% | 15 | 1.56% |
| Protein-Prospector | 10869 | 240 | 0 | 2.21% | 163 | 1.50% | 83 | 2.26% |
| Mango | 9497 | 0 | 0 | 0.00% | 78 | 0.82% | 108 | 1.96% |
| MS Annika | 7034 | 640 | 28 | 8.70% | 225 | 3.20% | 171 | 5.63% |
| MeroX | 1877 | 37 | 0 | 1.97% | 130 | 6.93% | 80 | 11.19% |
| pLink2 | 2946 | 91 | 25 | 2.24% | 229 | 7.77% | 71 | 10.18% |
| pLink3 (FDR) | 31941 | 646 | 8 | 2.00% | 3801 | 11.90% | 1695 | 17.21% |
| pLink3 (post-filter) | 25163 | 0 | 0 | 0.00% | 714 | 2.84% | 323 | 4.12% |
| XlinkX PD2.5 | 21543 | 12 | 529 | 2.51% | 7662 | 35.57% | 520 | 37.98% |
| XlinkX PD3.3 | 3416 | 133 | 4 | 3.78% | 109 | 3.19% | 45 | 4.51% |

**Supplemental Table S2.**
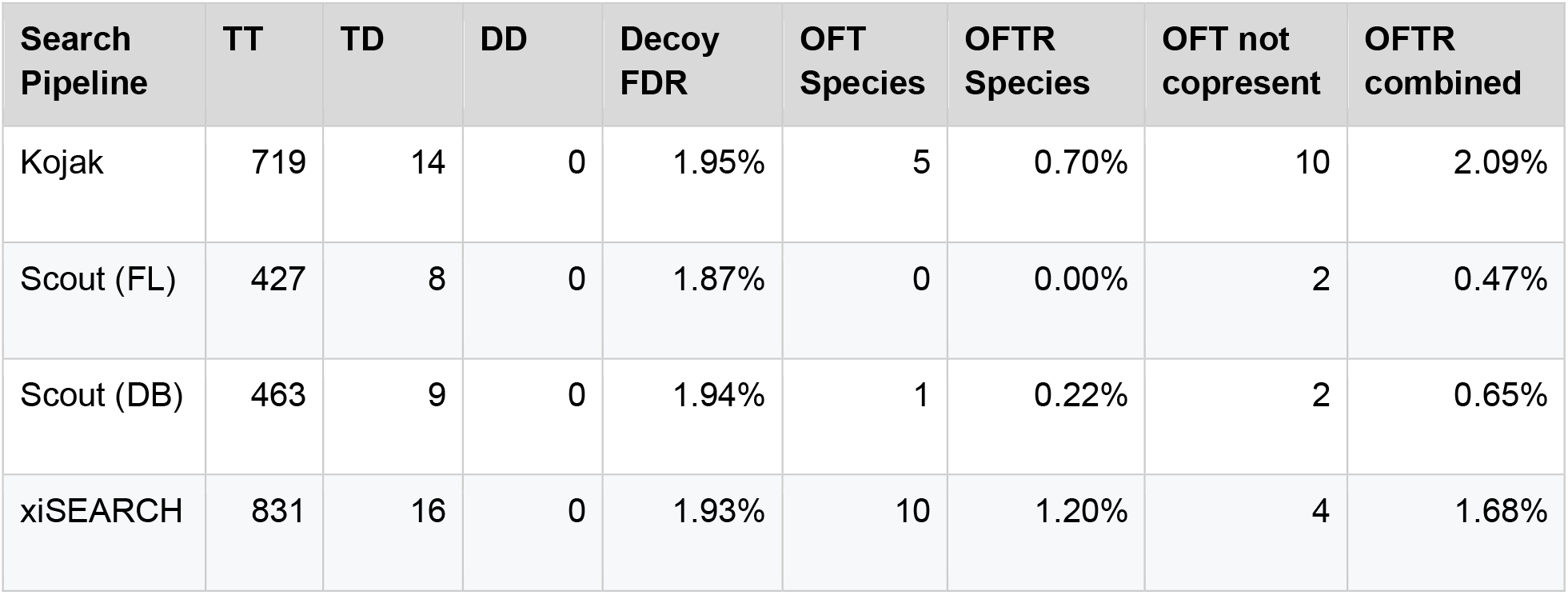
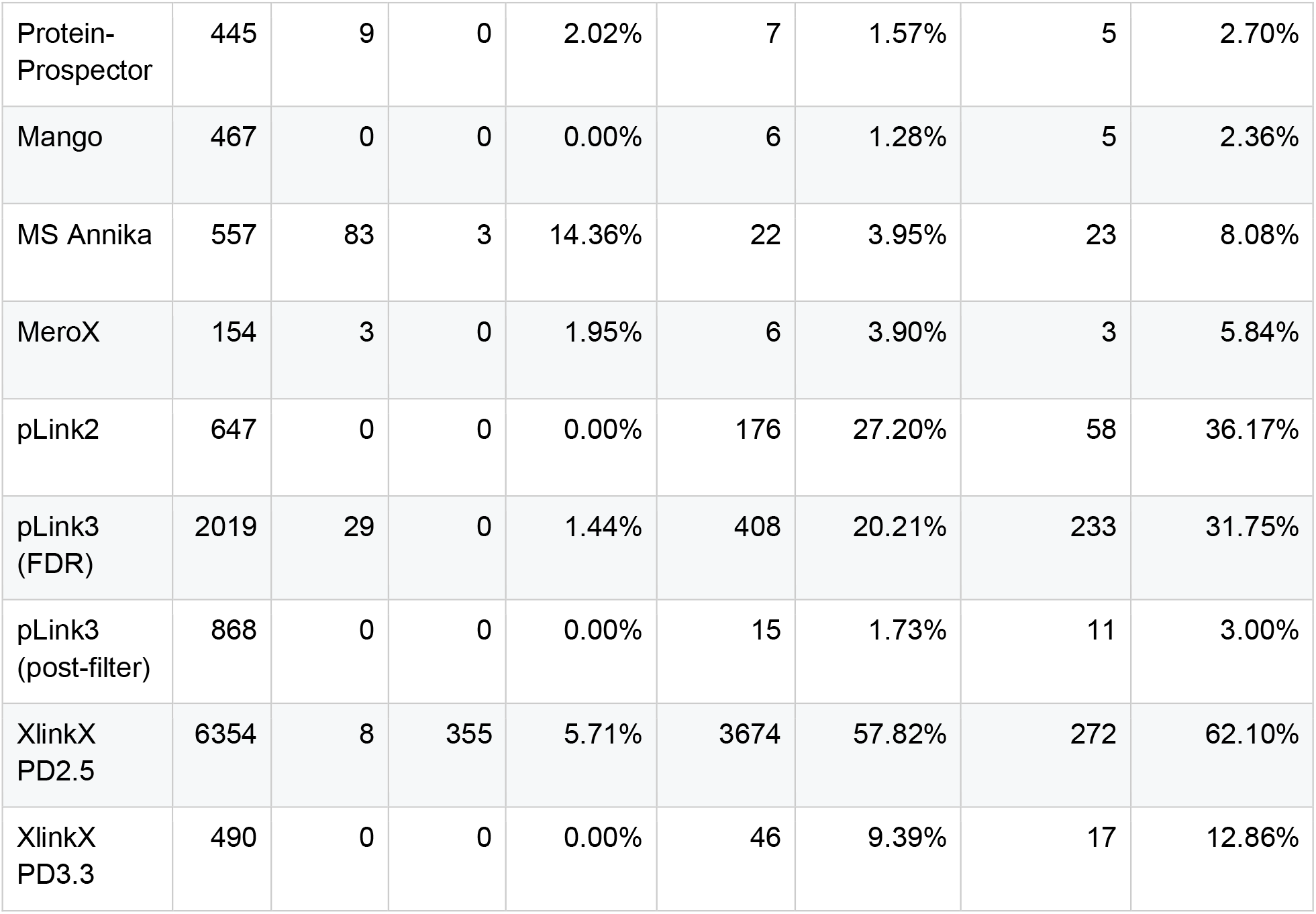
Single entrapment PPI level data per search engine.

**Supplemental Table S3.**
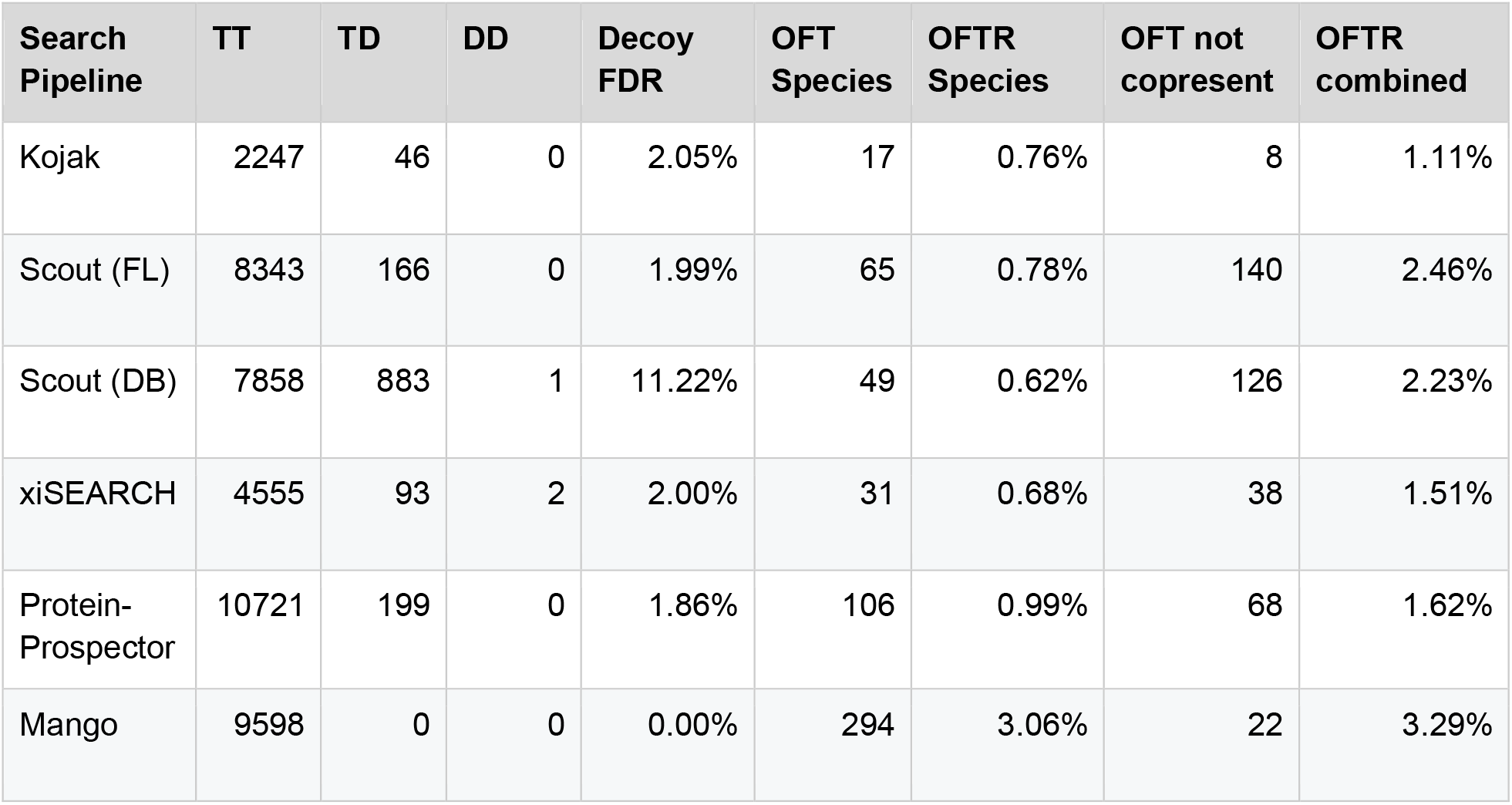
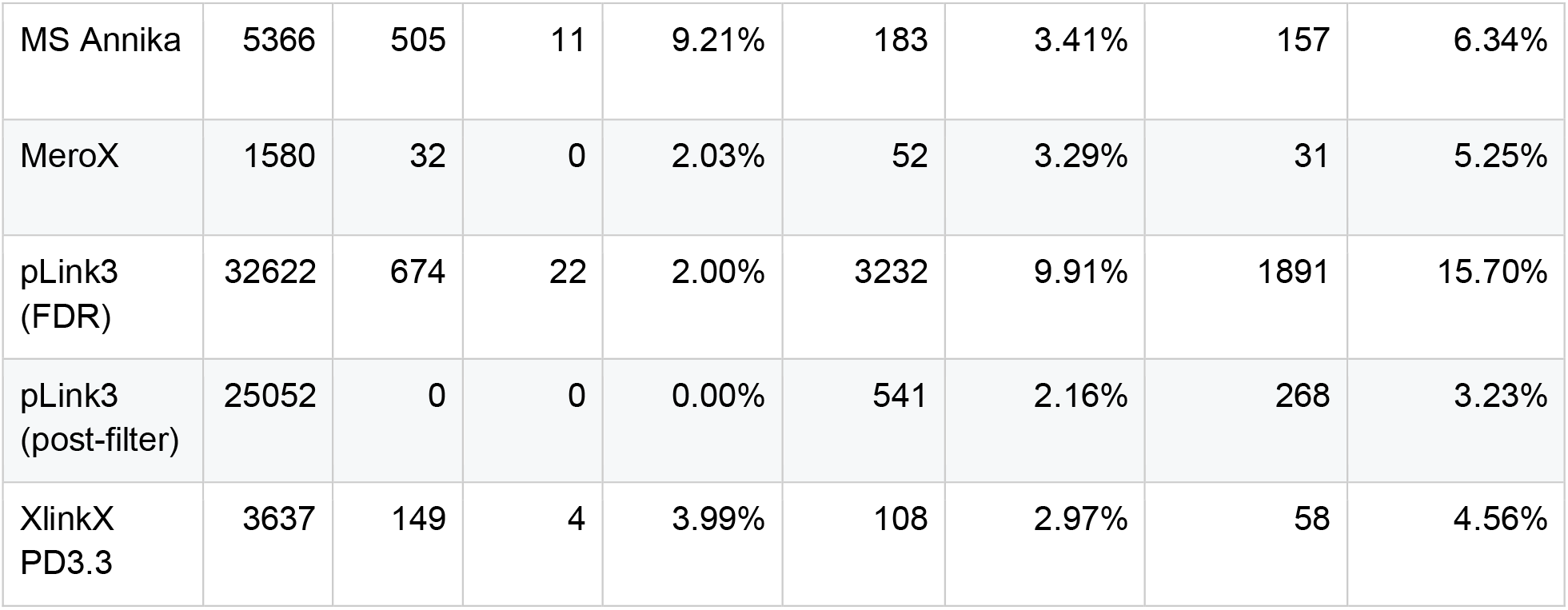
Dual entrapment CSM level data per search engine.

**Supplemental Table S4.** Dual entrapment PPI level data per search engine.

| Search Pipeline | TT | TD | DD | Decoy FDR | OFT Species | OFTR Species | OFT not copresent | OFTR combined |
| --- | --- | --- | --- | --- | --- | --- | --- | --- |
| Kojak | 114 | 2 | 0 | 1.75% | 1 | 0.88% | 1 | 1.75% |
| Scout (FL) | 506 | 10 | 0 | 1.98% | 3 | 0.59% | 2 | 0.99% |
| Scout (DB) | 565 | 12 | 0 | 2.12% | 6 | 1.06% | 6 | 2.12% |
| xiSEARCH | 838 | 16 | 0 | 1.91% | 13 | 1.55% | 8 | 2.51% |
| Protein-Prospector | 459 | 7 | 0 | 1.53% | 6 | 1.31% | 4 | 2.18% |
| Mango | 578 | 0 | 0 | 0.00% | 33 | 5.71% | 7 | 6.92% |
| MS Annika | 640 | 101 | 3 | 15.31% | 32 | 5.00% | 29 | 9.53% |
| MeroX | 157 | 3 | 0 | 1.91% | 6 | 3.82% | 3 | 5.73% |
| pLink3<br>(FDR) | 3140 | 53 | 2 | 1.62% | 733 | 23.34% | 402 | 36.15% |
| pLink3<br>(post-filter) | 996 | 0 | 0 | 0.00% | 29 | 2.91% | 10 | 3.92% |
| XlinkX<br>PD3.3 | 616 | 0 | 0 | 0.00% | 50 | 8.12% | 27 | 12.50% |

**Supplemental Table S5.**
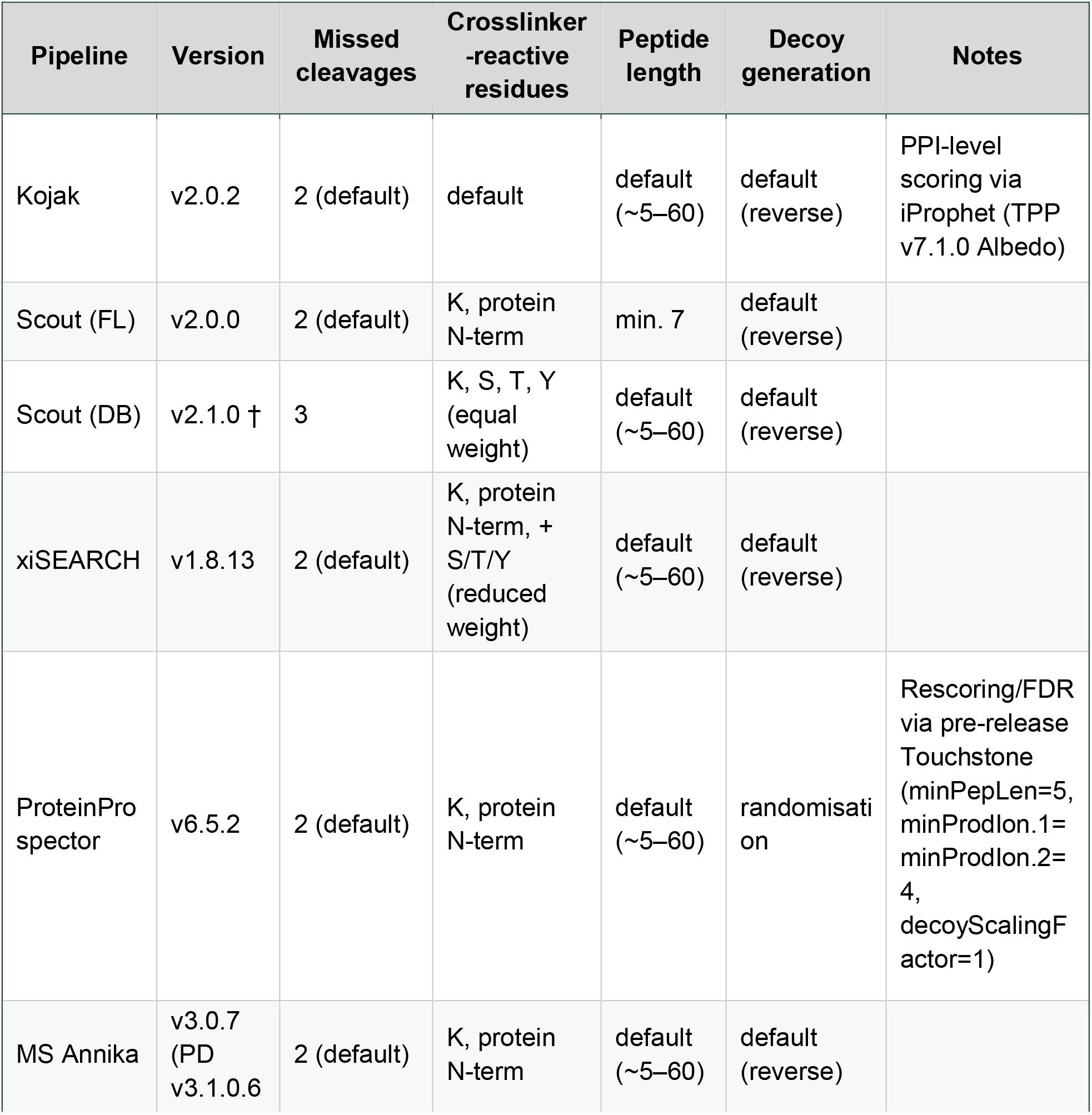
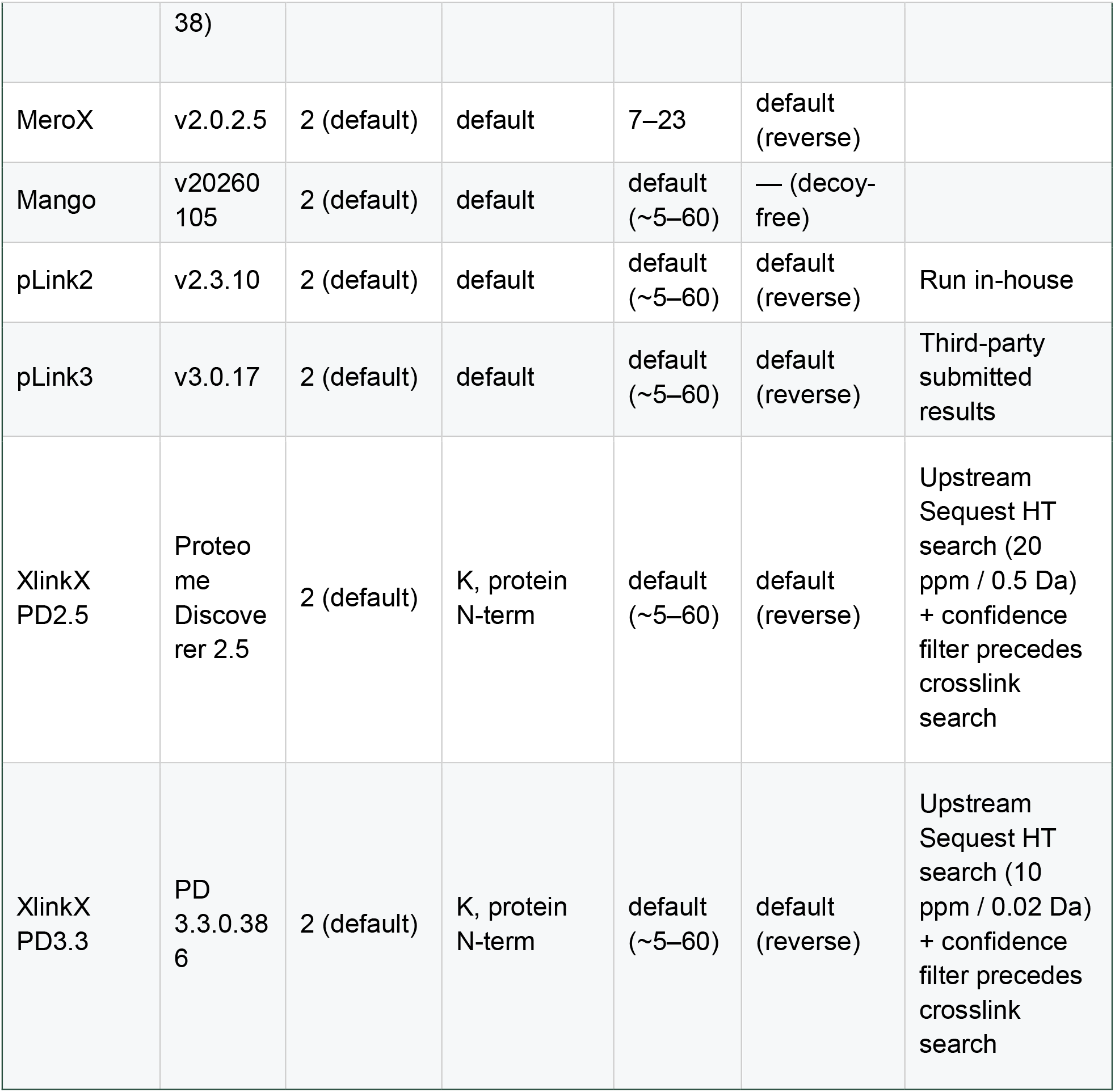
Search pipeline configurations. Crosslinker, protease, fixed/variable modifications, precursor/fragment tolerance (4 ppm / 8 ppm), and requested FDR (2%) were identical across all twelve pipelines and are omitted from the table; see above for the deviations summarised here.

## References

1. O’Reilly, F. J. & Rappsilber, J. Cross-linking mass spectrometry: methods and applications in structural, molecular and systems biology. Nat. Struct. Mol. Biol. 25, 1000–1008 (2018).

2. Walzthoeni, T. et al. False discovery rate estimation for cross-linked peptides identified by mass spectrometry. Nat. Methods 9, 901–903 (2012).

3. Fischer, L. & Rappsilber, J. Quirks of Error Estimation in Cross-Linking/Mass Spectrometry. Anal. Chem. 89, 3829–3833 (2017).

4. Iacobucci, C. et al. First community-wide, comparative cross-linking mass spectrometry study. Anal. Chem. 91, 6953–6961 (2019).

5. Leitner, A. et al. Toward increased reliability, transparency, and accessibility in Cross-linking mass spectrometry. Structure 28, 1259–1268 (2020).

6. Beveridge, R., Stadlmann, J., Penninger, J. M. & Mechtler, K. A synthetic peptide library for benchmarking crosslinking-mass spectrometry search engines for proteins and protein complexes. Nat. Commun. 11, 742 (2020).

7. Lenz, S. et al. Reliable identification of protein-protein interactions by crosslinking mass spectrometry. Nat. Commun. 12, 3564 (2021).

8. Matzinger, M. et al. Mimicked synthetic ribosomal protein complex for benchmarking crosslinking mass spectrometry workflows. Nat. Commun. 13, 3975 (2022).

9. Fischer, L. & Rappsilber, J. Rescuing error control in crosslinking mass spectrometry. Mol. Syst. Biol. 20, 1076–1084 (2024).

10. Clasen, M. A. et al. Proteome-scale recombinant standards and a robust high-speed search engine to advance cross-linking MS-based interactomics. Nat. Methods 21, 2327–2335 (2024).

11. Wen, B. et al. Assessment of false discovery rate control in tandem mass spectrometry analysis using entrapment. Nat. Methods 22, 1454–1463 (2025).

12. Bogdanow, B. et al. Redesigning error control in cross-linking mass spectrometry enables more robust and sensitive protein-protein interaction studies. Mol. Syst. Biol. 21, 90–106 (2025).

13. O’Reilly, F. J. et al. In-cell architecture of an actively transcribing-translating expressome. Science 369, 554–557 (2020).

14. Yang, B. et al. Identification of cross-linked peptides from complex samples. Nat. Methods 9, 904–906 (2012).

15. Hoopmann, M. R. et al. Kojak: efficient analysis of chemically cross-linked protein complexes. J. Proteome Res. 14, 2190–2198 (2015).

16. Hoopmann, M. R. et al. Improved analysis of cross-linking mass spectrometry data with Kojak 2.0, advanced by integration into the trans-proteomic pipeline. J. Proteome Res. 22, 647–655 (2023).

17. Shteynberg, D. et al. iProphet: multi-level integrative analysis of shotgun proteomic data improves peptide and protein identification rates and error estimates. Mol. Cell. Proteomics 10, M111.007690 (2011).

18. Mendes, M. L. et al. An integrated workflow for crosslinking mass spectrometry. Mol. Syst. Biol. 15, e8994 (2019).

19. Trnka, M. J., Baker, P. R., Robinson, P. J. J., Burlingame, A. L. & Chalkley, R. J. Matching cross-linked peptide spectra: only as good as the worse identification. Mol. Cell. Proteomics 13, 420–434 (2014).

20. Trnka, M. J. et al. Optimization of search and acquisition strategies for DSSO crosslinking mass spectrometry using Protein Prospector and Touchstone. bioRxiv (2026) doi:10.64898/2026.10.08.757733.

21. Birklbauer, M. J. et al. Proteome-wide non-cleavable crosslink identification with MS Annika 3.0 reveals the structure of the C. elegans Box C/D complex. Commun. Chem. 7, 300 (2024).

22. Götze, M. et al. Automated assignment of MS/MS cleavable cross-links in protein 3D-structure analysis. J. Am. Soc. Mass Spectrom. 26, 83–97 (2015).

23. Mohr, J. P., Perumalla, P., Chavez, J. D., Eng, J. K. & Bruce, J. E. Mango: A General Tool for Collision Induced Dissociation-Cleavable Cross-Linked Peptide Identification. Anal. Chem. 90, 6028–6034 (2018).

24. Chen, Z.-L. et al. A high-speed search engine pLink 2 with systematic evaluation for proteome-scale identification of cross-linked peptides. Nat. Commun. 10, 3404 (2019).

25. Liu, F., Lössl, P., Scheltema, R., Viner, R. & Heck, A. J. R. Optimized fragmentation schemes and data analysis strategies for proteome-wide cross-link identification. Nat. Commun. 8, 15473 (2017).

26. Heissel, S. et al. Fast and accurate disulfide bridge detection. Mol. Cell. Proteomics 23, 100759 (2024).

